# Elucidating the mechanism of cyclodextrins in the treatment of Niemann-Pick Disease Type C using crosslinked 2-hydroxypropyl-β-cyclodextrin

**DOI:** 10.1101/2020.07.31.230136

**Authors:** Dario Carradori, Hsintsung Chen, Beat Werner, Aagam Shah, Chiara Leonardi, Mattia Usuelli, Raffaele Mezzenga, Frances Platt, Jean-Christophe Leroux

**Affiliations:** Institute of Pharmaceutical Sciences, Department of Chemistry and Applied Biosciences, ETH Zürich, Zürich, Switzerland; Department of Pharmacology, University of Oxford, Oxford, United Kingdom; Center for MR-Research, University Children’s Hospital, Zürich, Switzerland; Institute of Neuroinformatics, ETH Zürich and University of Zürich, Zürich, Switzerland; Department of Health Sciences and Technology, ETH Zürich, Zürich, Switzerland

**Keywords:** Cholesterol, Crosslinking, Filipin, Magnetic Resonance Imaging-guided Low Intensity-Pulsed Focused Ultrasound, Niemann-Pick Disease Type C, 2-hydroxypropyl-β-cyclodextrin

## Abstract

Niemann-Pick Disease Type C (NPC) is a severe neurovisceral disorder that is pathophysiologically characterized by intracellular transport abnormalities leading to cytoplasmic accumulation of lipids such as cholesterol and multiple sphingolipids, including sphingosine. The compound 2-hydroxypropyl-β-cyclodextrin (HPβCD) is a compound with high cholesterol complexation capacity and is currently under clinical investigation for the treatment of NPC. However, due to its short blood half-life, high doses are required to produce a therapeutic effect. It has been reported in mice that HPβCD’s circulation time and efficacy can be improved by increasing its size *via* polymerization, but the biodegradable nature of these systems did not allow the contribution of the macromolecule to the activity to be determined. In this work, stable forms of polymerized HPβCD were generated (*via* epichlorohydrin crosslinking) to investigate their *in vitro* mechanisms of action and *in vivo* effects. Crosslinked CDs (8-312 kDa) displayed a 10-fold greater complexation capacity towards cholesterol than monomeric HPβCD but were taken up by cells to a lower extent (in a size-dependent fashion), resulting in an overall comparable *in vitro* effect on intracellular cholesterol accumulation that was dependent on cholesterol complexation. When tested *in vivo*, the crosslinked 19.3 kDa HPβCD exhibited a longer terminal half-life than the monomeric HPβCD. However, it did not increase the life span of *Npc1* mice, possibly due to reduced organ penetration and brain diffusion consequence of its large molecular weight. This could be circumvented by the application of magnetic resonance imaging-guided low intensity-pulsed focused ultrasound (MRIg-FUS), which increased the brain penetration of the CD. In conclusion, stable forms of polymerized HPβCD constitute valuable tools to elucidate CDs’ mechanism of action. Moreover, the use of MRIg-FUS to maximize CDs tissue penetration warrants further investigation, as it may be key to harnessing CDs full therapeutic potential in the treatment of NPC.

**Graphical abstract:** 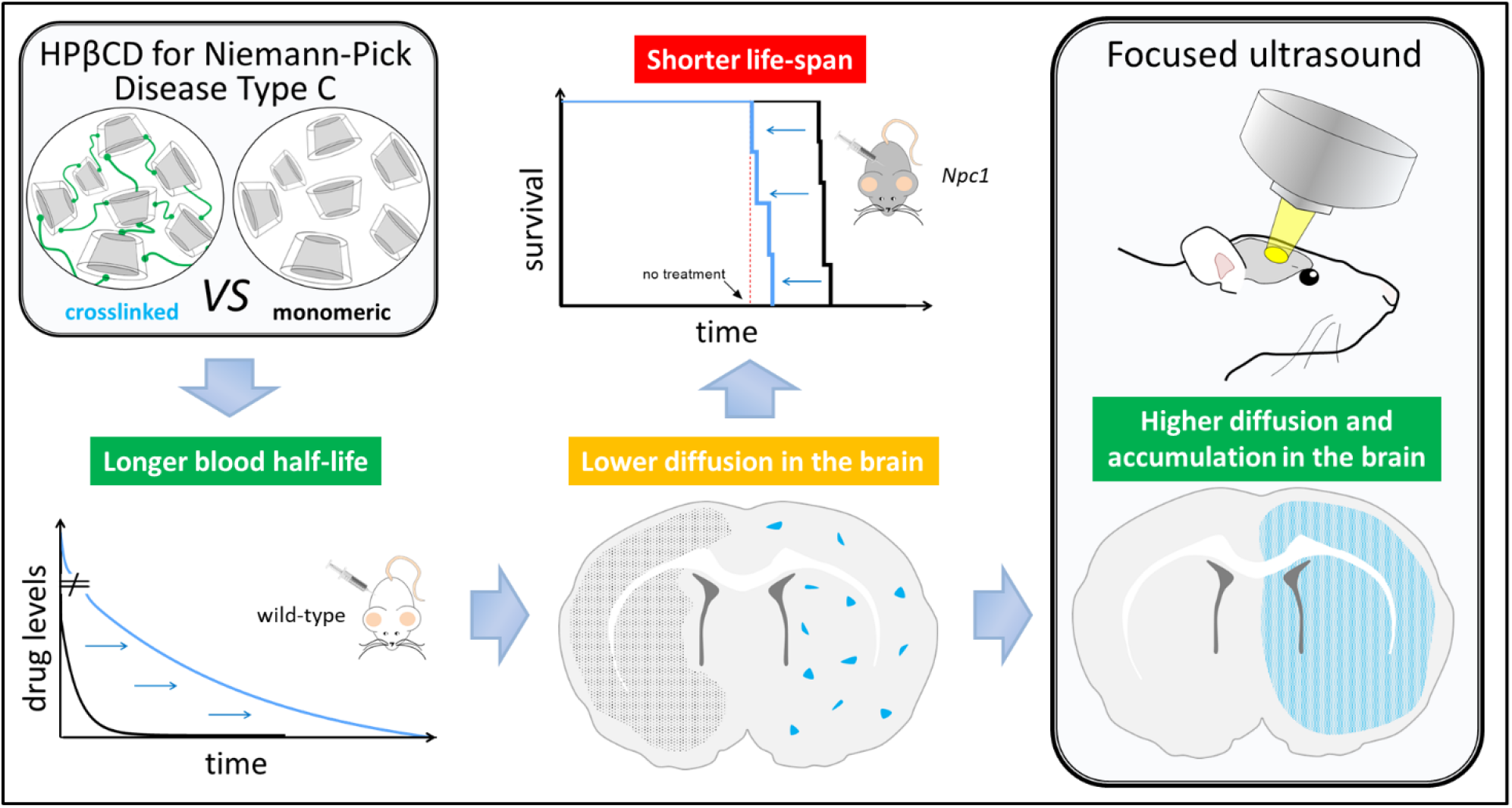

The 2-hydroxypropyl-β-cyclodextrin (HPβCD) is a well-established pharmaceutical excipient that can complex cholesterol and is currently under clinical investigation to treat Niemann-Pick Disease Type C (NPC). However, high doses of the drug are needed to achieve a therapeutic effect. Using stable and long circulating crosslinked HPβCDs, this study attempts to further understand the mechanisms behind CDs’ activity.

## 1. Introduction

Niemann-Pick Disease Type C (NPC) is a rare, inherited and prematurely fatal neurovisceral disorder attributed to the mutation of *Npc1* or *Npc2* genes (95% and 5% of the cases, respectively), which causes unique intracellular transport abnormalities of lipids and, subsequently, their accumulation in late endosomes/lysosomes (LE/LY) [1–4]. The clinical course of this lipid storage disorder is heterogeneous and depends on the age of onset [5–7]. In most cases, motor impairment and cognitive decline lead to premature death due to pneumonia and respiratory failure, or therapy-resistant epilepsy. At present, miglustat (Zavesca^®^) is the only drug approved for NPC treatment (in Europe, Brazil, South Korea, Canada and Japan). This iminosugar acts as a competitive inhibitor of the enzyme glucosylceramide synthase, which catalyses the first step in glycosphingolipid synthesis (*i.e*., the glycosylation of ceramide). The oral availability and the capacity to cross the blood-brain barrier (BBB) are the main advantages of this drug but the clinical benefits on NPC patients remains limited [8–11].

Different types of pharmacological approaches for NPC treatment are currently under investigation [12–17], and many of them focus on the regulation of lipid storage in the LE/LY [3]. The cyclic oligosaccharide 2-hydroxypropyl-β-cyclodextrin (HPβCD) is a well-established pharmaceutical excipient that has generated much interest as a potential treatment for NPC due to its high affinity for cholesterol and other NPC-relevant lipids [18–21]. The cyclodextrin (CD) drives a redistribution of the cholesterol from the LE/LY to the extracellular space but the mechanism behind it is not fully understood. *In vitro*, HPβCD has been shown to trigger the exocytosis of LE/LY content *via* a pathway that depends on the calcium channel MCOLN1 [22], and to promote lysosomal cholesterol exchange from the cellular plasma membrane to the serum lipoproteins [23], suggesting a dual action inside and outside the cell. HPβCD is in phase I/II [NCT03893071] and phase II/III [NCT03893071] clinical trials for NPC treatment under the name of Trappsol^®^ Cyclo^™^ and VTS-270, respectively. It is intravenously (i.v.) administered at high doses (1500 mg kg^−1^ or 2500 mg kg^−1^ over 8-9 h every two weeks) due to it short biological half-life (1.7 h [24]) and its low propensity to cross the BBB [18]. According to recent animal studies, the fraction of injected dose reaching the brain is very small [25], and relatively low levels (30-450 μg mL^−1^) have been detected in human cerebrospinal fluids 4 h after the beginning of the i.v. infusion. Consequently, strategies to increase HPβCD deposition in the brain are needed to enhance the efficacy of this treatment.

Recently, linear βCD-prodrugs with high molecular weight (*i.e*., an acid-labile βCD-based polyrotaxane with a molar mass of ~30 kDa [26] and a linear pH-degradable βCD-based polymer of ~33 kDa [27]) were shown to significantly increase the mean life span of NPC mice, while exhibiting a prolonged circulation time *vs*. HPβCD. These preliminary studies suggested that CD’s efficacy could be increased by reducing its systemic clearance. However, several mechanistic questions remain unanswered as neither of these studies investigated the impact of CD size on cholesterol complexation capacity, *in vitro* activity, and cellular uptake in a systematic way. Moreover, the contribution of the CD units (progressively released after administration) to the pharmacokinetics and biodistribution profile of the linear prodrugs was not evaluated and the polymeric CDs were not directly compared to their monomeric counterparts having identical molecular structures.

The molecular weight of CDs can be increased by several methods (*e.g*., by threading CDs to polymeric networks or by self-assembling host-guest inclusion complexes [28]), albeit crosslinking is one of the most straightforward approaches [29–32]. Epichlorohydrin (EPI) is the preferred crosslinker in polysaccharide chemistry [33] and EPI-derived crosslinked CDs have been found to complex hydrophobic molecules more than their native counterparts due to the presence of interstitial domains in the crosslinked polymer network [34]. One of the advantages of this approach is that EPI crosslinks are not biodegradable, allowing to study the impact of the molecular weight on CD activity, pharmacokinetics and biodistribution without the interference created by the release of the single units.

In this study, stable HPβCDs of high molecular weight (from 8 to 312 kDa) were synthetized *via* EPI crosslinking and characterized. Their cytotoxicity profile, *in vitro* activity (*i.e*., ability to decrease intracellular cholesterol accumulation) and cellular uptake were investigated on U18666A-induced NPC-fibroblasts. Furthermore, the pharmacokinetic and biodistribution profiles of a selected HPβCD were determined in healthy rats, and its therapeutic efficacy was assessed *in vivo* in a NPC animal model (*Npc1*^−/-^ mice) to gain insights on the mechanism of action of CDs in NPC. Finally, the use of magnetic resonance imaging-guided low intensity pulsed-focused ultrasound (MRIg-FUS) was explored as a mean to promote the brain penetration of CDs.

## 2. Results

### 2.1 Characterization of crosslinked HPβCDs

High molecular weight HPβCDs were synthesized by EPI crosslinking under alkaline conditions [35]. The molecular weight was varied by adjusting the reaction times, which were 6-fold shorter than those reported for βCD due to the higher reactivity of HPβCD. Five different crosslinked HPβCDs were prepared and characterized for their physicochemical properties (Table 1, Supplementary Figure 1S and 2S). The molar mass was found to increase rapidly when the reaction time exceeded 30 min, with the highest reproducibility obtained when the reaction time was set at 40 min, leading to the 19.3 kDa HPβCD. This crosslinked CD had a hydrodynamic diameter of ~6.5 nm which is slightly below the renal excretion threshold by filtration (*i.e*., 7 nm) [36].

**Table 1.** Crosslinked HPβCD characteristics. Two-way ANOVA with multiple comparisons among the parameters of the CDs within each column (Tukey’s post-hoc test). Mean ± SD (*n*=5 of independent batches, except *where *n*=4). Matching letters in superscript indicate a statistically significant difference (p < 0.05).

| Time (min) | MW (kDa) | D<sub>h</sub> (nm) | PDI | CD/EPI (%) | HP<sub>i</sub>/HP<sub>f</sub> (%) | Yield (%) |
| --- | --- | --- | --- | --- | --- | --- |
| 0 | 1.4 ± 0.6<sup>(A)</sup> | 2.5 ± 0.5 | 0.18<sup>(A)</sup> | 100 | 100 | / |
| 15 | 8.6 ± 3.8<sup>(A)</sup> | 5.7 ± 0.7 | 0.28<sup>(B)</sup> | 106 ± 11 | 101 ± 5.4* | 40.6 ± 17.8 |
| 30 | 9.3 ± 3.0<sup>(A)</sup> | 5.6 ± 1.2 | 0.29<sup>(B)</sup> | 86.0 ± 21.6 | 84.0 ± 16.3 | 41.9 ± 8.2 |
| 40 | 19.3 ± 4.4<sup>(A)</sup> | 6.6 ± 2.4 | 0.28<sup>(B)</sup> | 76.0 ± 17.3 | 77.0 ± 13.5 | 46.8 ± 10.3 |
| 45 | 46.0 ± 26.5<sup>(B)</sup> | 8.2 ± 2.4 | 0.35<sup>(C)</sup> | 75.0 ± 13.3 | 77.0 ± 26.8 | 47.6 ± 4.9 |
| 60 | 312 ± 216<sup>(B)</sup> | 21.0 ± 13.2 | 0.41<sup>(D)</sup> | 80.0 ± 8.3 | 81.0 ± 16.0 | 54.4 ± 16.0 |

MW: molecular weight, D<sub>h</sub>: hydrodynamic diameter, PDI: polydispersity index and HP<sub>i</sub>/HP<sub>f</sub>: initial/final hydroxypropyl groups ratios.

### 2.2 Cholesterol complexation capacity

The complexation capacity of crosslinked HPβCDs was determined by dissolving the latter in water in the presence of excess cholesterol. As shown in Figure 1, the crosslinking reaction substantially increased the ability of CDs to bind cholesterol (~10 fold), as previously reported for other lipophilic compounds [34]. The concentrations of solubilized cholesterol ranged between 2.7 and 7.5 μM for the monomeric HPβCD and reached 20 to 65 μM for crosslinked HPβCDs at an equivalent monomeric CD concentration. Differences were also observed among crosslinked HPβCDs, but they were small and did not follow a specific trend, possibly because of the relatively large polydispersity of the macromolecules (Table 1).

**Figure 1.**
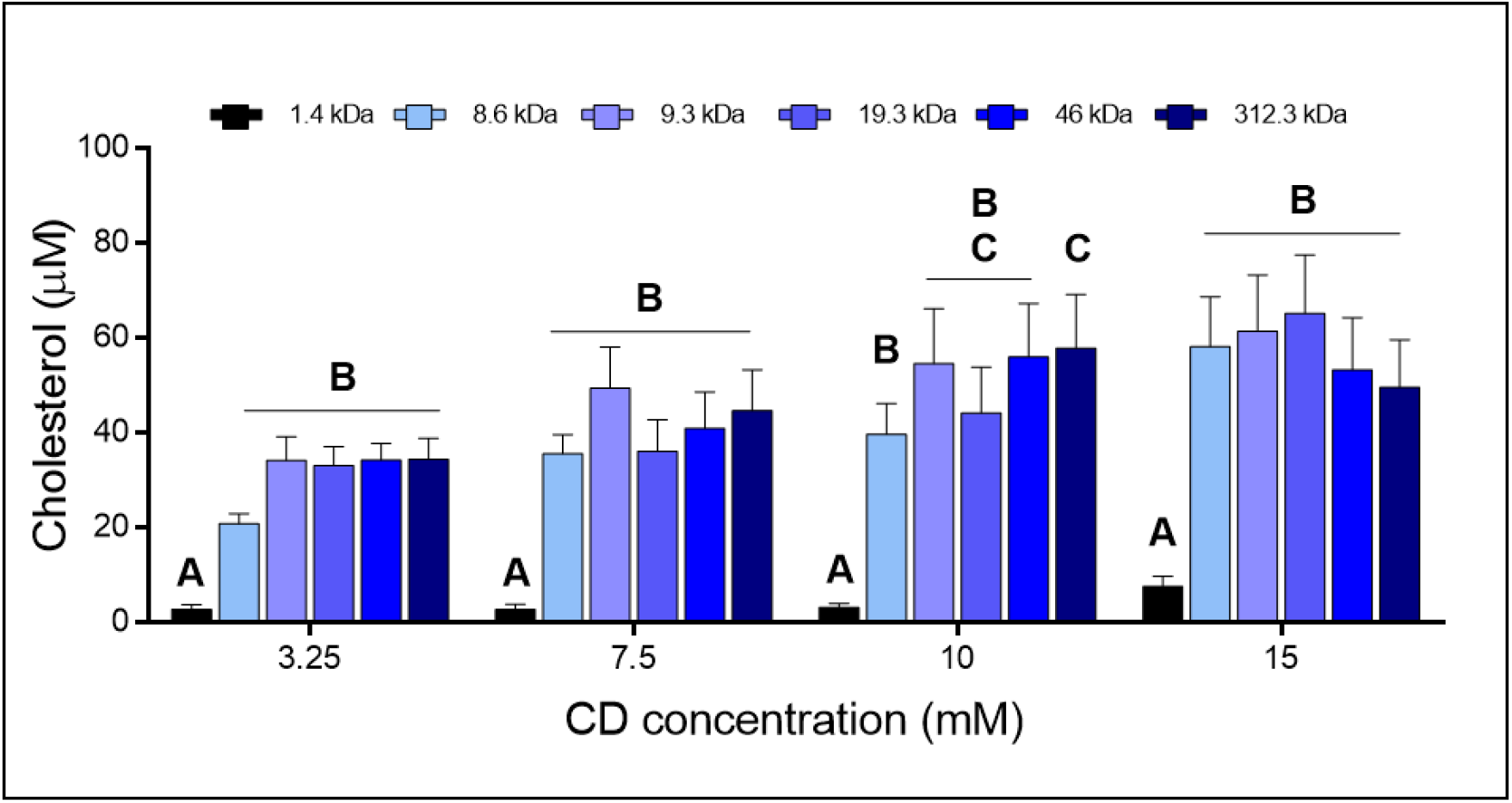
CD cholesterol complexation capacity at 3.25, 7.5, 10 and 15 mM. CD concentration is expressed in monomeric units. Two-way ANOVA with multiple comparisons (Tukey’s post-hoc test). Mean + SD (*n*=4). Different letters indicate a statistically significant difference (p < 0.05).

### 2.3 Impact on intracellular cholesterol accumulation

The NPC-phenotype was induced in mice fibroblasts (L929 cells) with a cholesterol transport inhibitor (U18666A) that binds NPC1 and was confirmed microscopically by filipin staining [37]. NPC-induced fibroblasts were incubated for 48 h with increasing concentrations of CDs in U18666A-free complete medium in order to evaluate the cytotoxicity profile (*i.e*. lactate dehydrogenase (LDH) release and viability) as well as the CD *in vitro* activity (*i.e*. decrease in intracellular cholesterol). NPC-like fibroblasts treated with all tested CD concentrations showed ≥80% viability *vs*. untreated cells, independently of CD molecular weight (Figure 2A, top panel). On the other hand, the LDH release progressively increased with CD concentrations (Figure 2A, bottom panel), likely due to the escalating deterioration of the plasma membrane integrity caused by CDs’ cholesterol extraction [38]. NPC-like fibroblasts treated with ≤5 mM HPβCDs released less than 10% of LDH compared to untreated cells while, at 10 mM, the LDH release increased up to ~10-20%. At 15 mM, monomeric HPβCD had a significantly higher impact (61%) than the other CDs (up to 25%), which could be explained by the higher cellular uptake of the monomeric *vs*. crosslinked HPβCDs (see paragraph 2.5). The cytotoxicity evaluation was also performed on normal fibroblasts with a similar outcome (Supplementary Figure 3S). Consequently, CD concentrations >10 mM were excluded from subsequent experiments to avoid significant membrane damage during cellular incubation with the CDs.

**Figure 2.**
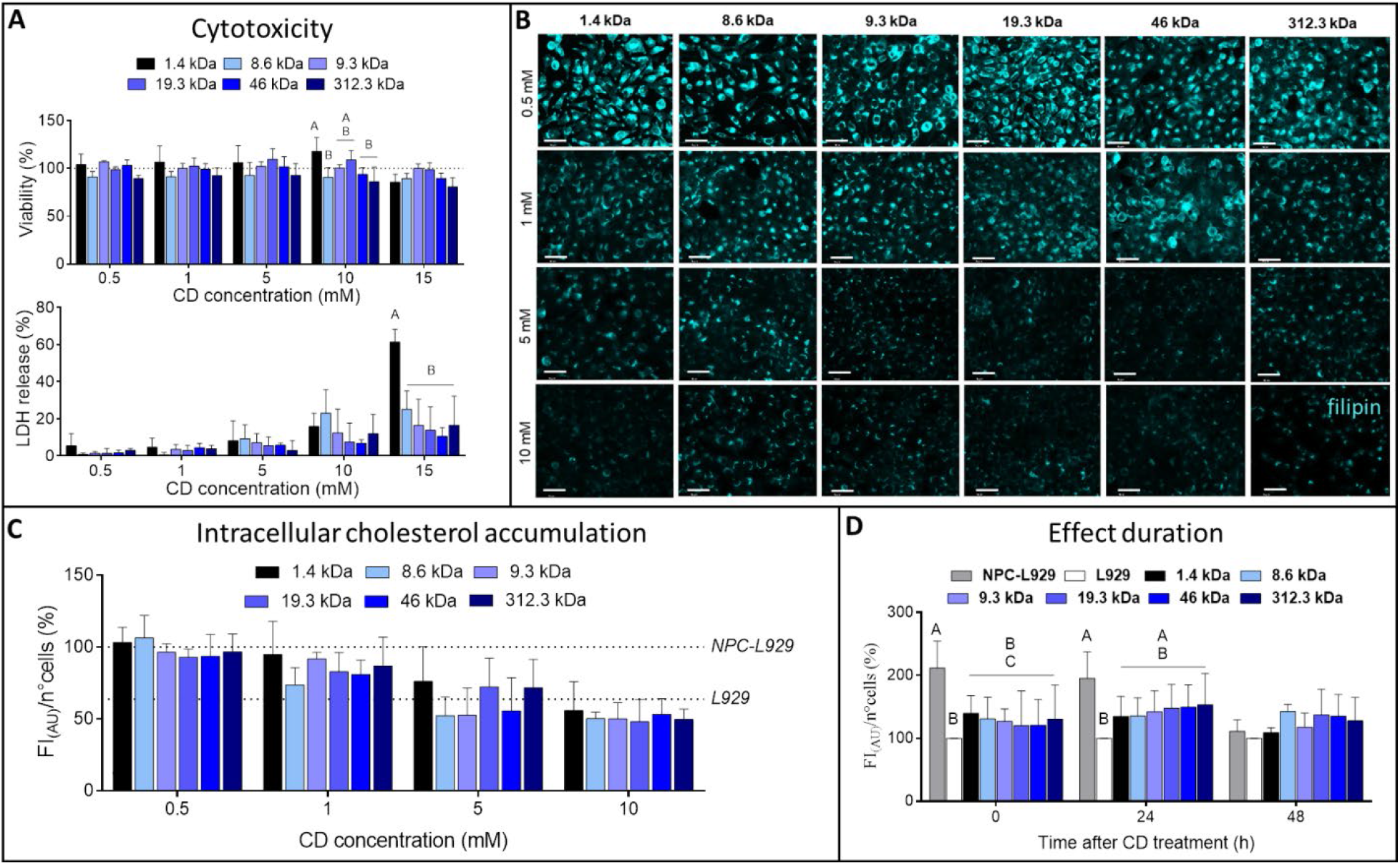
CD effect on intracellular cholesterol accumulation in NPC-like fibroblasts. NPC-L929 cells were treated with the different CDs (1.4 - 312.3 kDa) at increasing concentrations (0.5 – 15 mM). CD concentration is expressed in monomeric units. **A**) LDH release and cell viability (by MTS) after CD-treatment. **B**) Representative epifluorescence microscopy images of NPC-L929 cells after CD-treatment. Cyan corresponds to filipin fluorescence (*i.e*., intracellular cholesterol accumulation). Scale bar: 50 μm. **C**) Intracellular cholesterol quantification after CD-treatment. **D**) Duration of CD effect. Two-way ANOVA with multiple comparisons (Tukey’s post-hoc test). Mean + SD (*n*=3 for panel A, *n*=5 for B and C, *n*=4 for D). Different letters indicate a statistically significant difference (p < 0.05).

Filipin staining revealed a progressive reduction of cytoplasmic and perinuclear cholesterol clusters with increasing CD concentrations (Figure 2B). More specifically, no significant effect was observed at 0.5 mM while, above 1 mM, all CDs decreased the intracellular cholesterol in a concentration-dependent fashion (Figure 2C), with a trend towards a slightly higher activity for the crosslinked ones. The strongest effect was observed for concentrations ≥5 mM, where the intracellular cholesterol decreased down to the normal fibroblast’s level (*i.e*., 67%). Moreover, the shape of the cells became slightly rounded up when they were incubated for 48 h with 10 mM CDs (Supplementary Figure 3S), suggesting a possible morphology change not necessarily related to lower cell viability [39]. However, the differences in the *in vitro* efficacy were not statistically significant among the various CDs at any of the concentrations (Figure 2C). The 5 mM concentration was selected for subsequent experiments to ensure significant *in vitro* activity while excluding interferences due to cell morphology changes.

The impact of the CD size on the duration of the effect was investigated 24 and 48 h after CD-treatment (Figure 2D). The reduction of intracellular cholesterol induced by the CDs remained essentially unchanged for 24 h independently of their molecular weight. However, the NPC-phenotype was no longer detectable 48 h after CD-treatment since there was no significant difference between NPC-L929 and L929 values in the absence of CD treatment, probably due to cells’ recovery from the U18666A inhibition. Therefore, the crosslinking did not increase the duration of the CD effect compared to the monomeric HPβCD.

### 2.4 Kinetics and mechanism of CD cellular uptake

Monomeric (1.4 kDa), 19.3 and 312.3 kDa HPβCDs were selected to cover the small, medium and large size ranges, respectively, and were fluorescently labelled with rhodamine B (RhB) to study their cellular uptake. NPC-like fibroblasts were incubated with fluorescent CDs during 48 h and analyzed at specific time points to correlate the CD cellular uptake with the intracellular cholesterol reduction. The incubations with the fluorescent CDs were also performed at low temperature (4 °C) or in absence of ATP in order to determine whether uptake/activity were influenced by membrane fluidity and/or were energy dependent.

The kinetic studies over 48 h showed that the progressive uptake of CDs in the cells (Figure 3A, right panel) was accompanied by a concomitant reduction of the intracellular cholesterol accumulation (Figure 3A, left panel). At incubation times ≤4 h, CD cellular uptake and impact on intracellular cholesterol were comparable for both crosslinked CDs. The monomeric HPβCD led to a comparable reduction of intracellular cholesterol, despite its uptake being 6-fold higher than that of crosslinked HPβCDs. For longer incubation times (≥8 h), all CDs affected the intracellular cholesterol accumulation similarly, reducing it by −36% and −46% even though the monomeric HPβCD was taken up by cells 2- and 10-fold more than the 19.3 and 312.3 kDa HPβCDs, respectively. Representative images of the cells incubated with the CDs after different time-points are available in Supplementary Figure 4S. Furthermore, the intracellular localization of the fluorescent CDs was evaluated by scanning laser confocal microscopy. The orthogonal sectioning image analysis showed the progressive uptake of the CDs in the NPC-like fibroblasts over different time points (Supplementary Figure 5S), with a significant CD presence (monomeric and crosslinked) in the cytoplasm (Figure 3B, top panel) and at cholesterol accumulation sites after 48 h (Figure 3B, bottom panel).

**Figure 3.**
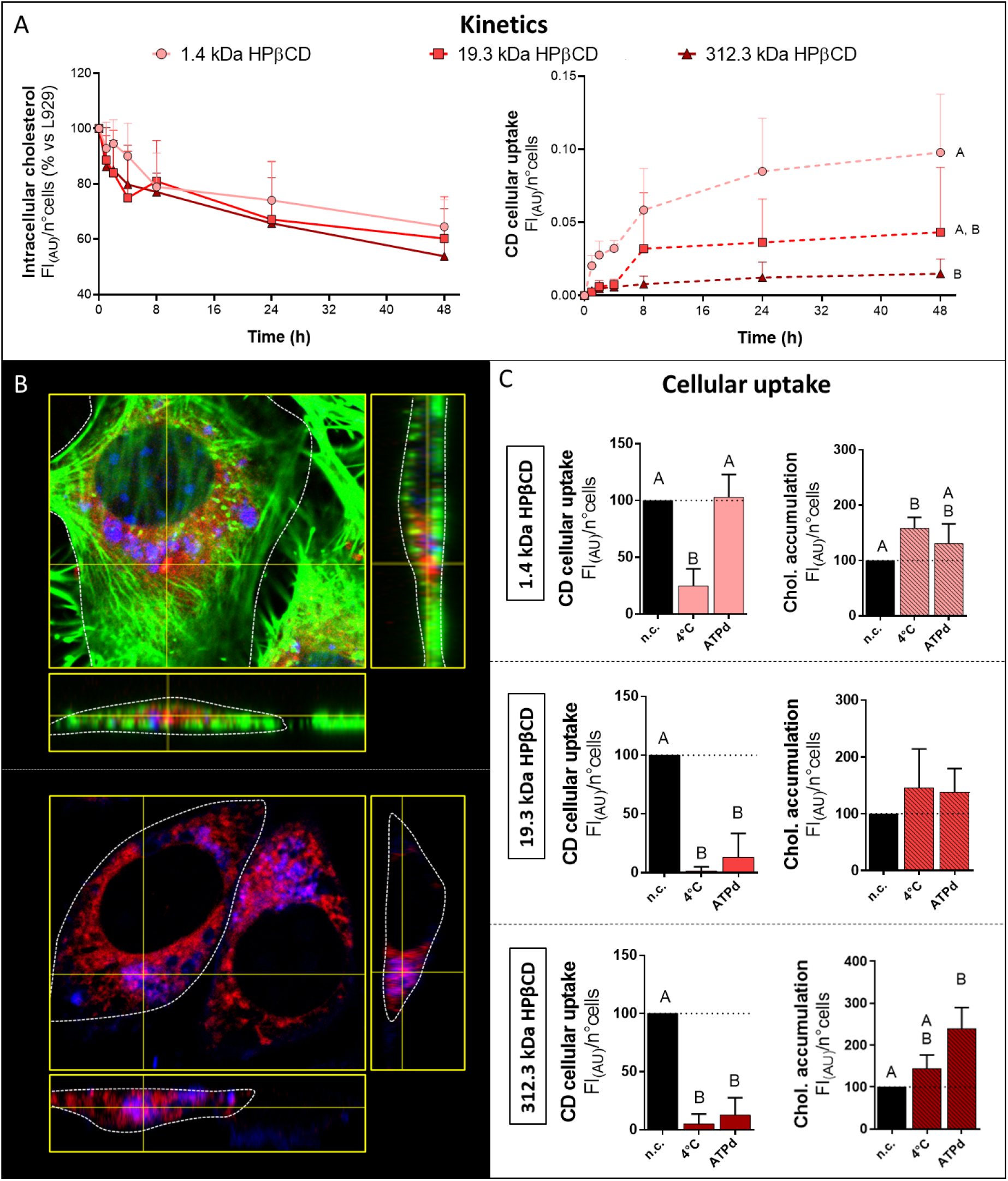
CD cellular uptake. **A**) Intracellular cholesterol (left panel) and CD cellular uptake (right panel) kinetics. Two-way ANOVA with multiple comparisons (Tukey’s post-hoc test). Mean + SD (*n*=5). **B**) Representative orthogonal sections of confocal microscopy images of CDs (monomeric HPβCD in this case) inside the cells (top panel) and colocalized with intracellular cholesterol (bottom panel). Blue corresponds to filipin (*i.e*., intracellular cholesterol), green corresponds to phalloidin (*i.e*., cytoskeleton), red corresponds to RhB (*i.e*., fluorescent CD), and violet corresponds to the colocalization between filipin and RhB. **C**) CD cellular uptake under normal conditions (n.c.), at low temperature (4 °C) or under ATP depletion (ATPd). One-way ANOVA with multiple comparisons (Tukey’s post-hoc test). Mean + SD (*n*=5). Different letters indicate a statistically significant difference (p < 0.05).

The CD uptake was modulated by temperature and ATP levels (Figure 3C). At 4 °C, the uptake of all fluorescent CDs decreased (−75% for monomeric, −98% for 19.3 kDa HPβCD and −95% for 312.3 kDa HPβCD), resulting in intracellular cholesterol levels 50-60% higher than those measured at 37 °C. On the other hand, the ATP depletion reduced the uptake of the crosslinked CDs exclusively (~ −80%), producing an intracellular cholesterol increase of 25% for 19.3 kDa HPβCD and 150% for 312.3 kDa HPβCD. The uptake of monomeric HPβCD was not impacted, though a small but significant increase (~ 25%) of intracellular cholesterol was observed.

### 2.5 Modulation of the CD in vitro activity by the CD inclusion site

The importance of having free CD inclusion sites to achieve efficacious intracellular cholesterol complexation in NPC-like fibroblasts was investigated. Using CDs that were pre-complexed with cholesterol, the variation of intracellular cholesterol accumulation was quantified microscopically by filipin staining (Figure 4). Pre-complexation with cholesterol significantly affected the monomeric HPβCD function, resulting in a 75% activity loss. Interestingly, when the exogenous cholesterol and the monomeric HPβCD were added separately, the effect on intracellular cholesterol complexation was preserved, indicating the importance of having a free inclusion site in the oligosaccharide structure. On the other hand, the activity of the crosslinked HPβCDs was not significantly reduced by the pre-complexation in comparison to adding the CDs and cholesterol separately. Therefore, the crosslinking seemed to counteract the activity loss observed in the pre-complexed monomeric HPβCD, probably *via* the interstitial inclusion sites of the polymeric network.

**Figure 4.**
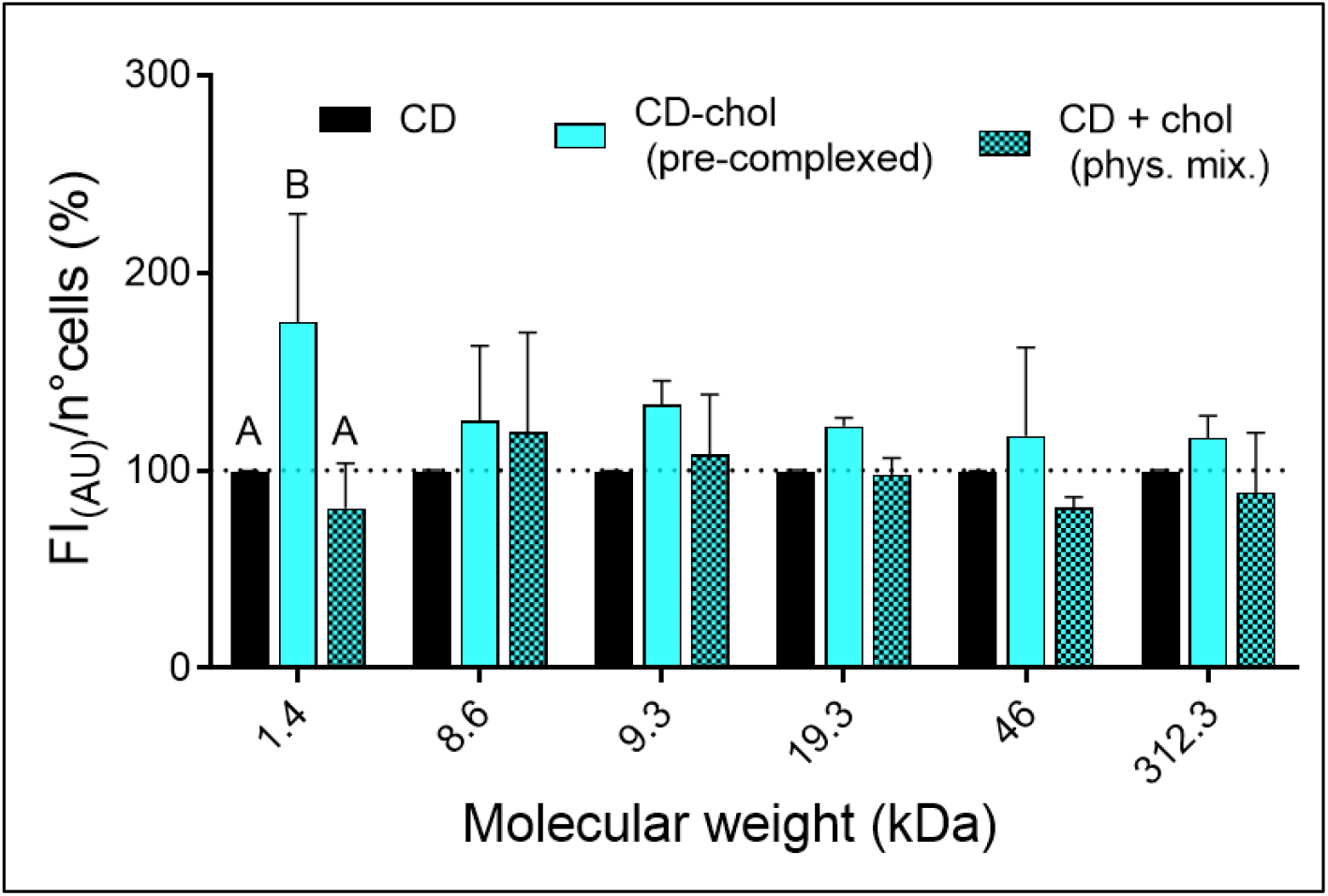
Role of CD inclusion site after pre-complexation with cholesterol on *in vitro* activity. Two-way ANOVA with multiple comparisons (Tukey’s post-hoc test). Mean + SD (*n*=3).

### 2.6 Pharmacokinetic study

Despite having a similar *in vitro* activity to the other crosslinked CDs (Figures 1–4), the 19.3-kDa HPβCD was selected for the pharmacokinetics and efficacy experiments based on its favorable size, which should contribute to a longer half-life *in vivo* (*i.e*., 2.5 times larger than native HPβCD but close to the renal filtration threshold).

Following i.v. injection in wild-type rats, the 19.3-kDa HPβCD displayed a prolonged circulation time compared to the monomeric HPβCD (Table 2, Figure 5A). The terminal half-life (t_1/2_) and mean residence time (MRT) increased from 2.6 to 3.8 h and 0.2 to 2.3 h, respectively, while the clearance (CL) was reduced by 40%, likely reflecting a slower elimination from the body. Moreover, the apparent volume of distribution (Vd) decreased from 0.086 to 0.076 L kg^−1^, reflecting the increased retention in the blood pool.

**Figure 5.**
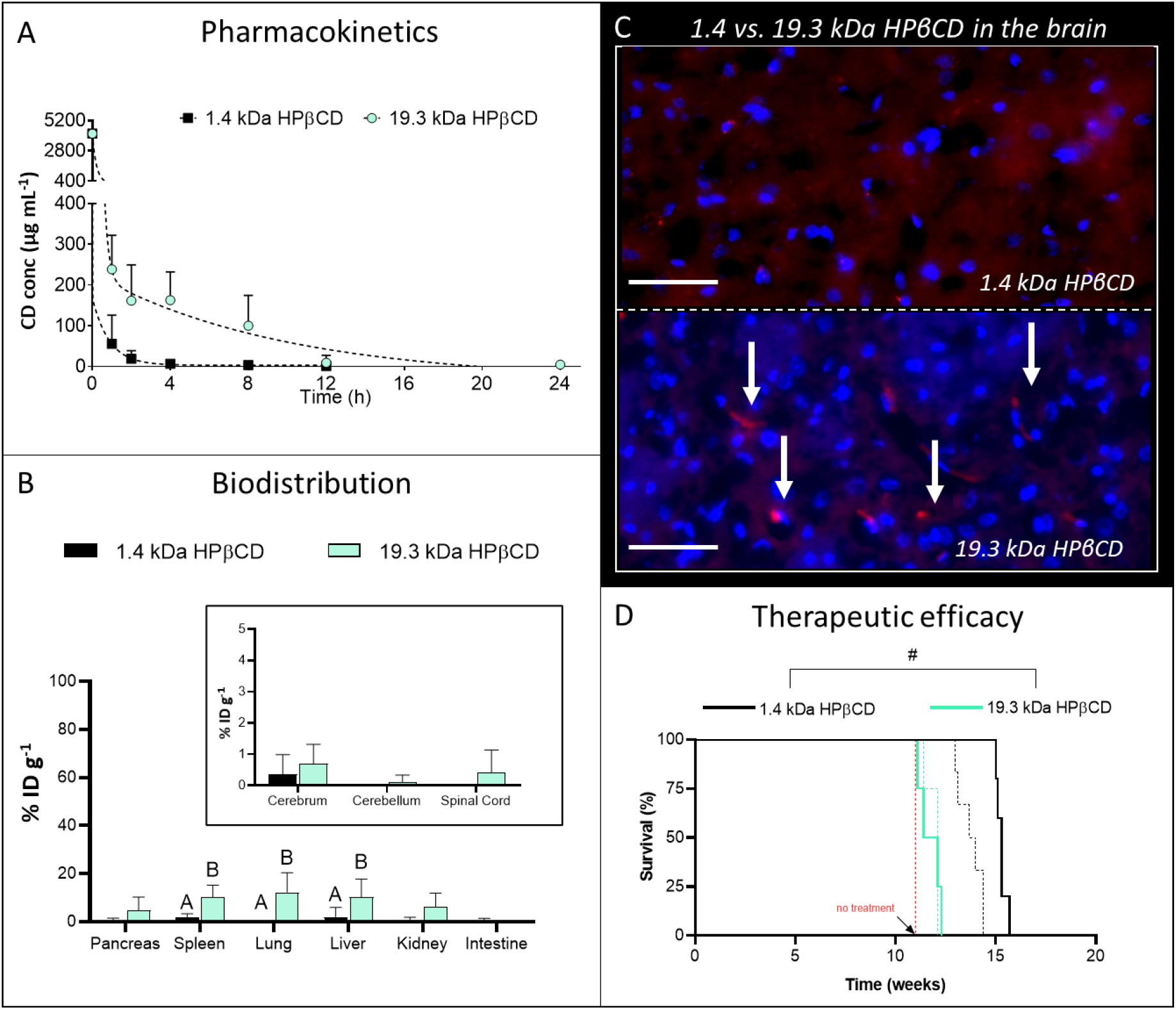
CD pharmacological properties. **A**) CD pharmacokinetics in rat plasma at different time-points. **B**) CD biodistribution in rat main organs 12 h after CD i.v. administration. Two-way ANOVA with multiple comparisons (Tukey’s post-test). Mean + SD (*n*=6 for cerebrum, cerebellum and spinal cord, *n*=7 for the other organs). Different letters indicate a statistically significant difference (p < 0.05). **C**) Representative pictures epifluorescence microscopy images of monomeric *vs*. crosslinked CD diffusion in the brain parenchyma. Blue corresponds to Hoechst (*i.e*., cell nuclei), red corresponds to RhB (*i.e*., fluorescent CD), scale bar: 50 μm. **D**) CD therapeutic efficacy in P7 Npc1 mice after CD i.p. injection at 1333 mg kg^−1^ (dotted line) or 4000 mg kg^−1^ (continuous line). Kaplan-Meier survival plot with Log-rank test (Mantel-Cox) between the CDs (*n*=4 for 19.3 kDa HPβCD at both doses, *n*=6 for 1.4 kDa HPβCD at 1333 mg kg^−1^ and *n*=5 for 1.4 HPβCD kDa at 4000 mg kg^−1^). The symbol # indicates a statistically significant difference (p < 0.05).

**Table 2.** Pharmacokinetic parameters of i.v. injected monomeric and crosslinked CDs.

| Parameters | Unit | 1.4 kDa HPβCD | 19.3 kDa HPβCD |
| --- | --- | --- | --- |
| $t_{1/2}$ | h | 2.62 | 3.78 |
| $C_{\max}$ | $\mu\text{g mL}^{-1}$ | 4160 | 4160 |
| $\text{AUC}_{0-t}$ | $(\mu\text{g mL}^{-1}) \text{ h}$ | 2200 | 3550 |
| CL | $\text{mg h}^{-1}$ | 0.023 | 0.014 |
| $V_d$ | L kg<sup>-1</sup> | 0.086 | 0.076 |
| MRT | h | 0.2 | 2.3 |

$t_{1/2}$ : terminal half-life, $C_{\max}$ : maximum serum concentration, $\text{AUC}_{0-t}$ : area under the curve from 0 to infinity, CL: clearance, $V_d$ : volume of distribution, MRT: mean residence time.

Twelve hours after CD injection, 19.3 kDa HPβCD was mainly found in lung, liver and spleen (~ 10% ID g^−1^), followed by the kidney and the pancreas (~ 5% ID g^−1^), while the monomeric HPβCD could only be recovered in small amounts in the liver and spleen (~ 2% ID g^−1^) (Figure 5B). The distribution in the central nervous system was generally extremely low (≤0.5 % ID g^−1^) for both CDs. Traces of the 19.3 kDa HPβCD were found in the spinal cord (0.4 % ID g^−1^). Since the concentration of both types of CDs seemed similar in the brain, their penetration in the parenchyma was compared. The 19.3 kDa HPβCD was confined in small clusters while the 1.4 kDa HPβCD diffused widely and homogeneously in the tissue (Figure 5C).

In order to test the *in vivo* activity, monomeric (1.4 kDa) and crosslinked (19.3 kDa) HPβCD were intraperitoneally (i.p.) injected in Npc1 mice at two different doses (1333 and 4000 mg kg^−1^). The selected dose was based on previous reports [40–42] showing efficacy after single dosing. In the literature [43], it was reported that in this model, mice with no treatment had a life span of about 11 weeks. The administration of the crosslinked HPβCD did not appear to increase survival, resulting in a lifespan of ~ 11-12 weeks. Indeed, the therapeutic effect was significantly inferior compared to that of the monomeric HPβCD at the two tested doses, *i.e*. 13.8 and 15. 3 weeks at 1333 and 4000 mg kg^−1^, respectively (Figure 5D).

### 2.7 Magnetic resonance imaging-guided low intensity-pulsed focused ultrasound

The limited *in vivo* therapeutic activity of the 19.3 kDa HPβCD was attributed to its increased retention in the blood pool (Table 2) and lower diffusion in the organs in comparison to the monomeric HPβCD (Figure 5B), in particular in the brain (Figure 5). In a pilot study in wild type rats, magnetic resonance imaging-guided low intensity-pulsed focused ultrasound (MRIg-FUS) was applied to transiently open the BBB and increase the accumulation and diffusion of the monomeric and 19.3 kDa HPβCD in the brain parenchyma.

Extravasation of gadolinium (Figure 6A) and Evan’s blue (Figure 6B) indicated an increased BBB permeability in the site of MRIg-FUS application. Therefore, 19.3 kDa HPβCD was i.v. injected during the FUS sequence and its brain accumulation was evaluated 8 h after CD administration. The macromolecule showed a stronger and wider diffusion compared to the control regions (*i.e*., ultrasound-untreated areas) (Figure 6C). Furthermore, the CD was found in the proximity of astrocytes and oligodendrocytes (Figure 6D), suggesting a diffusion towards the neural cells rather than an accumulation on the epithelial cells of the blood vessels. The combination of i.v. injected monomeric CD with MRIg-FUS produced similar results (Supplementary Figure 6S).

**Figure 6.**
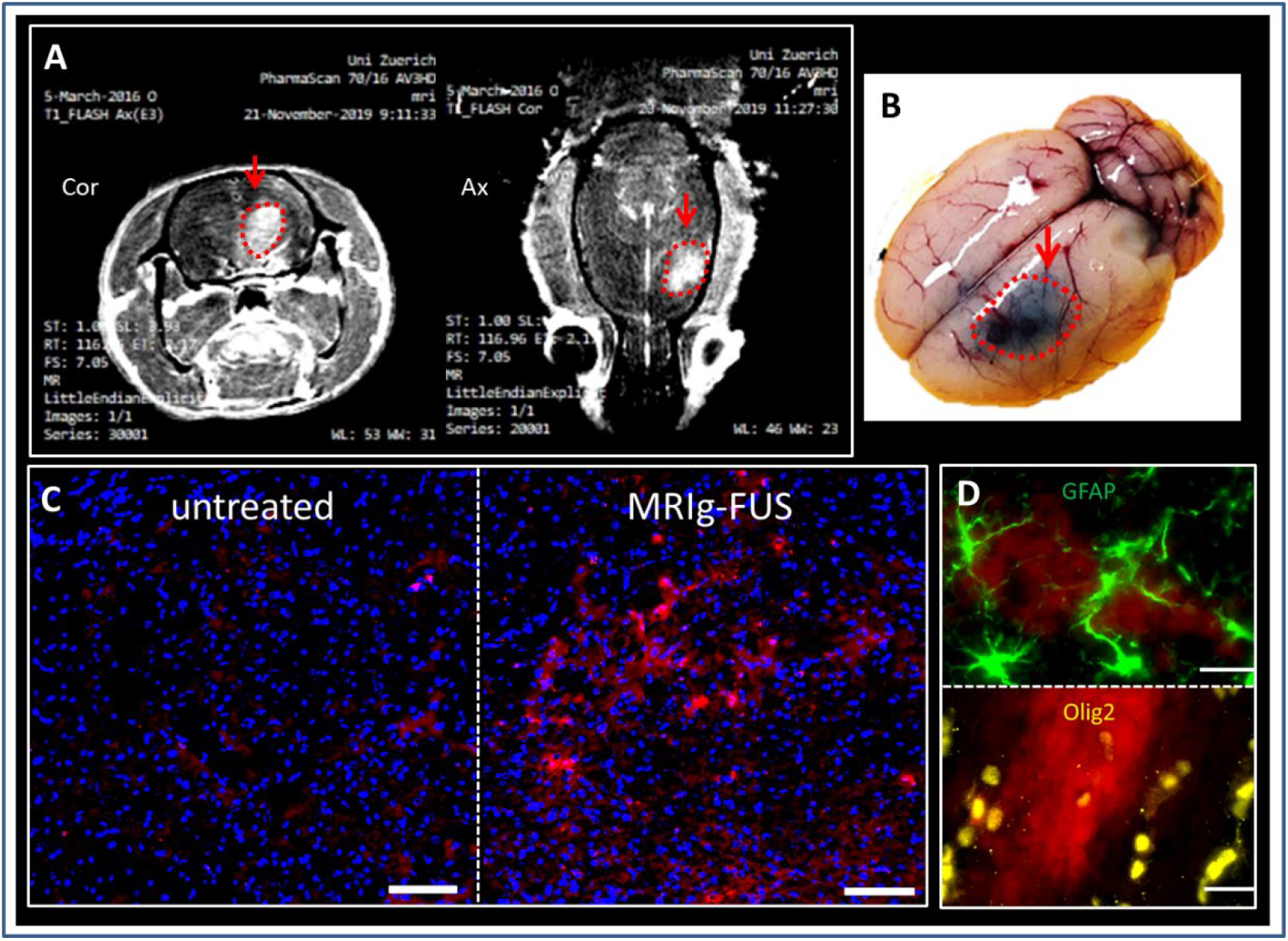
Brain diffusion of crosslinked CD after MRIg-FUS. **A**) T1-weighted MRI of gadolinium extravasation (clear grey) after MRIg-FUS. **B**) Full brain picture of Evan’s blue extravasation (dark blue) after MRIg-FUS. **C**) Representative microscopy images showing the diffusion of 19.3 HPβCD in untreated (left) and MRIg-FUS treated (right) brain regions. Blue corresponds to Hoechst (*i.e*., cell nuclei), red corresponds to RhB (*i.e*., fluorescent CD), scale bar: 100 μm. **D**) Representative microscopy images showing the localization of 19.3 HPβCD compared to the astrocytes and oligodendrocytes. Green corresponds to GFAP (*i.e*., astrocytes), yellow corresponds to Olig2 (*i.e*., oligodendrocytes), red corresponds to RhB (*i.e*., fluorescent CD), Scale bar: 20 μm.

## 3. Discussion

NPC is a devastating neurovisceral disorder with a profound impact on the patient’s life, imposing tremendous limitations in basic physiological and social needs. The progression of the disease leads NPC patients to inexorably lose their autonomy, while financial costs, emotional stress, and logistical complexities increase [1–7]. To date, miglustat is the only approved NPC treatment but its clinical benefit is limited. Several pharmacological approaches are currently under clinical trial, including HPβCD, adrabetadex [NCT03643562], lithium carbonate [NCT03201627], N-acetyl-L-leucine [NCT03759639], arimoclomol [NCT02612129, NCT04316637], vorinostat [NCT02124083] and pluripotent stem cells [NCT03883750]) [44]. HPβCD is one of the most studied molecules for NPC treatment and is being tested in several clinical trials, six of them are currently active (source: clinicaltrials.gov). However, there is no consensus on its mechanism of action [45] nor on the need to cross the BBB to display efficacy [46]. Recent reports indicate that increasing the molecular weight of CDs could represent a useful approach to improve the therapeutic outcome by prolonging the circulation time of these drugs [19]. On the other hand, increasing the molecular weight can also decrease the amount of drug reaching the brain due to the BBB [47]. Previous studies relied on macromolecular CD systems [26, 27], which were designed to gradually release monomeric βCD. However, they did not distinguish between the released unit and the linear prodrug, complicating interpretations of the impact of the molecular weight on the efficacy. In this work, stable crosslinked HPβCD with non-biodegradable linkers were developed as a tool to gain insight into the mechanism of action of CDs.

The simple modulation of the reaction time during EPI crosslinking produced different molecular weight HPβCDs. The presence of interstitial (“extra”) inclusion sites in the branched polymeric network made these compounds significantly more capable of cholesterol complexation in solution (up to 10-fold) than the monomeric HPβCD. The higher complexation capacity, however, did not result in a higher *in vitro* efficacy in NPC-phenotype fibroblasts. Crosslinked HPβCDs were taken up by the cells less efficiently (from 2- to 10-fold) than monomeric HPβCD. Consequently, it is possible that the lower cell uptake of the crosslinked HPβCDs counteracted their high cholesterol complexation capacity, thwarting its *in vitro* effect. Pre-complexation with cholesterol had been previously characterized physicochemically [48, 49] and used to reduce the toxicity of high CD concentrations by decreasing CD-induced cholesterol extraction from plasma membrane. However, its impact on the CD-mediated intracellular cholesterol accumulation had not been investigated *in vitro* [50]. By pre-loading the CDs with cholesterol at a ratio sufficient to saturate the cavities of the CD units (but not those created by the crosslinks), it was possible to establish that cholesterol complexation is crucial for the *in vitro* activity. This central role in the mechanism of action is in line with a recent publication where βCDs were less efficient *in vitro* when their inclusion site was stably occupied by a polymeric chain [51]. Indeed, our data showed that pre-filling CD inclusion sites generally abolished the *in vitro* efficacy of monomeric HPβCD, whereas the effect was much less pronounced with the crosslinked CDs, which were still capable of capturing cholesterol in the void spaces created by the crosslinks.

Our data also showed that the cellular uptake was key for the activity of either types of CDs. The crosslinked CDs were mainly taken up *via* an energy-dependent process (likely endocytosis), which could be expected considering that their large molecular weight would hamper their diffusion through the cell membrane [52]. In contrast, the fact that monomeric HPβCD uptake was only reduced at 4 °C and not under energy depleting conditions would support the theory that a diffusive process might be involved [53]. Indeed, it is known that the rigidity of the cell membrane increases at low temperature, which might impair interactions with the CDs as already observed with other compounds [54]. These findings confirm previous data from Vacca *et al*., [21] suggesting that cellular uptake is a prerequisite for CDs to significantly decrease intracellular cholesterol accumulation. The authors showed that an important part of the CD activity takes place inside the cell by interacting with a pathway dependent on the calcium channel MCOLN1 [21]. Thus, the kinetics and mechanism of cellular internalization of the CD can dictate its efficacy.

The 19.3 kDa HPβCD was selected for the in *vivo* studies, mainly because its hydrodynamic diameter would make this polymer large enough to increase circulation time, while still allowing its glomerular filtration [55]. The pharmacokinetics profile obtained for the monomeric HPβCD was in line with previously published data [26, 27], showing rapid and nonlinear clearance. On the contrary, the 19.3 kDa HPβCD exhibited longer plasma half-life and decreased clearance, being completely cleared from the bloodstream 24 h after injection. This is in contrast with previous reports on macromolecular βCD-based systems with higher molecular weight (30 kDa [26] and 33 kDa [27]), which were still clearly detectable after 24 and 48 h, respectively. After 12 h, the crosslinked HPβCD was found in small amounts mainly in the organs of the mononuclear system (liver and spleen) and lungs, while monomeric HPβCD was mostly cleared from the body. The concentrations of CD detected in the brain (cerebrum and cerebellum) were extremely low and not significantly different between monomeric and crosslinked HPβCDs, as previously reported for the biodegradable 30 kDa linear polymer [26]. The fact that after 12 h the monomeric HPβCD was still detectable in the brain is probably due to its slow brain clearance [56]. The prolonged blood circulation time did not lead to a high penetration of the crosslinked CD into the brain. Here, at the same % ID g^−1^, the monomeric HPβCD showed a wider and more homogeneous diffusion compared to the crosslinked HPβCD that could be attributed to the size difference.

The *in vivo* activity of the crosslinked HPβCD was assessed in a Npc1^−/-^ mouse model in which the single i.p. injection of an HPβCD dose in P7 old mice had been reported to result in a significant prolonged life span [40–42]. Despite having a comparable *in vitro* activity on NPC-like fibroblasts and longer circulation time, the crosslinked CD was significantly less effective than the monomeric counterpart at the tested doses, which was the opposite of previously reported findings with the other macromolecular βCD-based systems [26, 27]. Aside from the fact that the animal model, dosing regimen, administration route and nature of the monomeric CD (*e.g*., HPβCD *vs*. βCD) were not the same as in those previous studies, the discrepancy could be explained by the fact that in our system, monomeric CDs were not released from the macromolecular structure due to the non-biodegradability of the crosslinking [34, 35]. It is possible that given the lower volume of distribution of the macromolecule and its limited diffusion into the organs’ parenchyma, the 19.3 kDa HPβCD may not be able to diffuse deep enough in tissues to exert its activity. It is also possible that since crosslinked CDs are taken up by an active endocytotic process, any cells that are unable to passively internalize these larger structures in sufficient amounts would not contribute to the overall *in vivo* activity.

The limited brain delivery and penetration was addressed employing MRIg-FUS, which has been shown to enhance the deposition of different drugs in various organs [57–59] while being well tolerated by animals [60–63] and humans [64, 65]. Moreover, MRIg-FUS showed therapeutic efficacy *per se* on phosphorylated Tau and amyloid plaques [66, 67]. In our exploratory study, brain accumulation and diffusion of the 19.3 kDa HPβCD were substantially improved after only one application of FUS. Therefore, the combination of MRIg-FUS with the i.v. administration of CDs may represent a new promising strategy for the treatment of NPC. Indeed, it would allow both monomeric and crosslinked HPβCDs to achieve higher doses in the brain, answering the critical question about the importance of brain penetration in the activity of CDs.

To conclude, stable crosslinked CDs proved to be valuable tools to study the mechanism of action of CDs, revealing that both cholesterol complexation and cellular uptake are important determinants of CDs’ capacity to lower the intracellular accumulation of cholesterol. By increasing the hydrodynamic ratio of HPβCD, it was possible to increase its circulation time. However, the non-biodegradable crosslinks impaired tissue penetration and reduced its activity. This could be partially overcome with MRIg-FUS, with which promising results were obtained promoting the penetration of high molecular weight CDs in the brain parenchyma. In view of the technology’s increasing availability in hospitals worldwide, MRIg-FUS should be further investigated for its potential implementation in NPC patients receiving CDs or other treatments [44]. In view of previously published work, these data further suggest that sustained release formulations of CDs, for instance in the form of a biodegradable CD-based implant, could potentially simplify the dosing regimen while potentially improving the treatment efficacy. On the other hand, MRIg-FUS promotes the penetration of high molecular weight CDs in the brain parenchyma and should be further investigated. Indeed, this technology is becoming more easily accessible in the hospitals worldwide and could therefore be implemented in the near future in NPC patients receiving CDs or other treatments [44].

## 4. Experimental section

### 4.1 Animals

Jugular vein-catheterized adult female Sprague Dawley rats were purchased from Janvier Labs (Le Genest Saint Isle, France). Rats were maintained on a constant 12:12 h light:dark cycle while water and food were provided ad libitum. Pharmacokinetic and biodistribution studies were performed in accordance with procedures and protocols approved by the cantonal veterinary authorities (Kantonales Veterinäramt Zürich, license ZH218/17). The animals were euthanized with an overdose of xylazine and ketamine.

The 9 females and 10 males Npc1 mice (BALBc/NPC^nih^, BALB/cNctr-*Npc1*^m1N^/J) were generated from heterozygous mice (*Npc1*^+/-^) breeding stock obtained from The Jackson Laboratory (The Jackson Laboratory, Charles River, United Kingdom) [54]. Mice were maintained on a constant 12:12 h light:dark cycle with food and water available ad libitum. The experiments with these mice were conducted using protocols approved by the United Kingdom home Office Animal Scientific Procedures Acts 1986 (P8088558D).

### 4.2 Materials

HPβCD was purchased from CycloLab (Budapest, Hungary). Sodium hydroxide 98.5-100% pellets (NaOH), 15 mm Ø No.1 coverslips and hydrochloric acid (HCl) 37% were purchased from VWR International (Radnor, PA). Filipin III complex *Streptomyces filipinensis* 75% (filipin), acetone 99%, maleic acid ≥ 99%, paraformaldehyde 95% (PFA), epichlorohydrin 99% (EPI), triphenylphosphine (Ph3P) ≥ 95%, sodium azide (NaN3) ≥ 99.5%, sodium methoxide (NaOCH3) ≥ 97%, dichloromethane anhydrous (DCM) ≥ 99.8%, palladium on carbon (PdC) 10 wt. %, hydrazine hydrate 98%, rhodamine b isothiocyanate (RhB) mixed isomers, 2-deoxy-D-glucose ≥ 98%, and iodine (I2) 99.9%, Evan’s blue 75% were purchased from Merck KGaA (Darmstadt, Germany). Spectra/Por^®^3 dialysis membranes MWCO 3500 were purchased from Spectrum Labs (Rancho Dominguez, CA). Syringe filter units Filtropur S 0.45 μm were purchased from Sarstedt (Nümbrecht, Germany). Pierce^™^ LDH Cytotoxicity Assay Kit and Amplex^™^ Red Cholesterol Assay Kit were purchased from Invitrogen (Waltham, MA). 96-well black microplates were purchased from Greiner bio-one (Kremsmünster, Austria). Deuterium oxide was obtained from Cambridge Isotope Laboratories (Tewksbury, MA). NCTC clone 929 cells (L929, mouse fibroblasts) were from the American Type Culture Collection (Manassas, VA). CellTiter 96^®^ Aqueous One Solution Cell Proliferation Assay Kit was purchased from Promega AG (Dübendorf, Switzerland). Micro BCA^™^ Protein Assay Kit, Gibco^®^ Fetal Bovine Serum, Hoechst, rabbit anti-Olig2 polyclonal antibody, rabbit anti-GFAP polyclonal antibody, Alexa goat anti-rabbit Alexa 488 secondary antibody, Gibco^®^ DMEM-high glucose-HEPES-no phenol red, Gibco^®^ Trypsin-EDTA-phenol red (0.25%), Gibco^®^ MEM-GlutaMAX^™^ and Gibco^®^ Penicillin-Streptomycin (10,000 U mL^−1^) were from ThermoFisher Scientific (Waltham,MA). Dimethylformamide (DMF) extra pure was purchased from Fisher Chemical (Hampton, NH). Methanol (CH3OH) 99.8% and pyridine 99.5% were purchased from Acros Organics (Geel, Belgium). 1,2-Distearoyl-sn-Glycero-3-Phosphocholine (DSPC) was from Avanti Polar Lipids (Alabaster, AL). DSPC-N-[Methoxy(Polyethylene glycol)-2000] (DSPE-PEG2000) and DSPC-N-[methoxy(polyethylene glycol)-5000] (DSPE-PEG5000) were purchased from Corden Pharma (Liestal, Switzerland). Syringes for the production of the microbubble were purchased from Tyco Healthcare (Mansfield, MA).

### 4.3 Methods

#### 4.3.1 Synthesis of crosslinked HPβCDs

Crosslinked HPβCDs were prepared adapting the protocol described by Renard *et al*. [35]. Briefly, 3 g of HPβCD were dissolved in 7 mL of NaOH solution (35% w/w) at 25 °C. The solution was then heated up to 30 °C and 1.72 mL of EPI were added while stirring. After 15, 30, 40, 45 or 60 min, the reaction was stopped by adding 10 mL of acetone. The aqueous phase containing the CD was collected *via* separating funnel. The solution was neutralized using HCl and dialyzed against deionized water for 3 days using a 3.5-kDa cut-off membrane. The ratios EPI/HPβCD and initial/final hydroxypropyl groups in the crosslinked HPβCDs were obtained by an ^1^H NMR AV400 400 MHz spectrometer (Bruker, Billerica, MA) (Supplementary Figure 1S) [68]. ***Fluorescently labelled CDs*** - The fluorescent CDs were prepared following the method described in Malanga *et al*. [69] in 3 steps: azidation, amination and rhodamine B isothiocyanate (RhB) labelling. Azidation was achieved by dissolving 16 mg of Ph3P (0.06 mmol) and 16 mg of I2 (0.06 mmol) in 20 mL of extra pure DMF under stirring at a temperature below 40 °C. Then, 1.3 mmol of CD (monomeric, 19.3 kDa or 312.3 kDa HPβCD) were added to the solution and the temperature was increased up to 50 °C and stirred for 1 h. The temperature of the solution was decreased down to 30 °C, and 100 mL of CH3OH with 0.1 g of NaOCH3 (1.85 mmol) were added. The solution was kept under stirring for 30 min before removing CH3OH with a rotary evaporator at 40 °C for 1 h at 2 kPa. Afterwards, 2 mL of DMF with 8 mg of NaN3 (0.12 mmol) were added and the solution was heated up to 80 °C and stirred for 2 h. DMF was removed by rotary evaporation at 80 °C for 2 h at 2 kPa and by high vacuum overnight. Subsequently, 20 mL of water were added to the dry material and filtered by glass filtration. The filtrate was extracted by adding 40 mL of DCM. The aqueous phase was frozen overnight and lyophilized for 2 days (yield around 90%). Amination was achieved by dissolving the lyophilized compound in 15 mL of water. Then, 0.18 g of PdC, previously suspended in 2 mL of water, and 1 mL of hydrazine hydrate (0.02 mol) were added to the solution and heated to reflux (at 105 °C) for 1 h. The solution was cooled down and PdC was removed by glass filtration (3 times) and centrifugation (4000 x *g* for 30 min). The supernatant was collected, and the pH was adjusted to 4-5. The solution was frozen overnight and lyophilized for 2 days (yield around 90%). RhB labelling was achieved by dissolving 1 g of the lyophilized compound and 6 mg of RhB in 10 mL of pyridine at 65 °C and stirred the mixture for 20 h. Pyridine was removed from the solution by rotary evaporator at 60 °C for ~ 5 h at 2 kPa and by high vacuum overnight. Then, 50 mL of water were added to dissolve the powder that was extracted 3 times with DCM and then dialyzed with 2- and 3.5-kDa cut-off membranes, for monomeric and crosslinked CDs, respectively, during 24 h. The solution was frozen overnight and lyophilized for 2 days (yield around 94%).

#### 4.3.2 Static and dynamic light scattering measurements

The molecular weight of crosslinked HPβCDs was measured using a 3D-LS Spectrometer (LS Instruments, Fribourg, Switzerland). The range of investigated concentrations was between 1 and 10 wt%. The refractive index increment of dn/dc = 1.26 x 10^−4^ mL mg^−1^ was empirically obtained with a RFM 340 refractometer (Bellingham & Stanley Ltd, Tunbridge Wells, United Kingdom) and data were analyzed by Debye Plot [70, 71] (Supplementary Figure 2S). The hydrodynamic diameter and polydispersity index were measured using a Malvern Zetasizer Nano ZS (Malvern Instruments, Herrenberg, Germany) with a crosslinked HPβCD concentration of 5 wt%. All samples were prepared in nanopure water and filtered through a 0.45-μm syringe filter.

#### 4.3.3 Solubility studies of cholesterol-CD complexes

The cholesterol complexation capacity of the crosslinked HPβCDs was measured following the method described by dos Santos *et al*. [57]. Briefly, an excess amount of cholesterol (1.5 mg) was added to a 10-mL aqueous solution containing increasing concentrations (0–15 mM) of monomeric or crosslinked HPβCDs and stirred for 48 h at 55 °C. The samples were then centrifuged (4000 x *g*, 4 °C, 2 h) and the supernatants were collected and filtered through a 0.45-μm syringe filter. The concentration of solubilized cholesterol (*i.e*., the cholesterol complexed by the CDs) was measured in each sample with the Amplex^™^ Red Cholesterol Assay Kit following the provider’s instructions.

#### 4.3.4 U18666A-induced NPC-like fibroblasts

L929 cells (*i.e*., mouse fibroblasts) were treated to develop the NPC-phenotype by following a protocol described by Lange *et al*. [58]. Briefly, fibroblasts were cultured at 37 °C, 5% CO2 in MEM GlutaMAX^™^ supplemented with 10% fetal bovine serum and 1% penicillin/streptomycin (complete medium). The cells were seeded at 2.5 x 10^4^ mL^−1^ and incubated with 4 μM U18666A cholesterol transport inhibitor for 72 h. The medium containing U18666A was removed and replaced by specific media depending on the experiment’s aim.

#### 4.3.5 Cytotoxicity experiments

L929 cells were seeded at 2.5 x 10^4^ mL^−1^ in a 96-well plate (200 μL/well) and treated to develop the NPC phenotype. The NPC-inducing medium was removed and replaced by complete medium with increasing concentrations of CDs (0.5, 1, 5, 10 and 15 mM) for 48 h. At the end of the treatment, the medium of each condition was collected to quantify lactic dehydrogenase (LDH) by Pierce^™^ LDH Cytotoxicity Assay Kit while the cells were analyzed by CellTiter 96^®^ Aqueous One Solution Cell Proliferation Assay Kit (*i.e*., MTS), following each providers’ instructions. The values obtained from CD-untreated cells were considered as 0% LDH release and 100% cell viability for LDH and MTS assays, respectively.

#### 4.3.6 In vitro intracellular cholesterol modulation

L929 cells were seeded at 2.5 x 10^4^ mL^−1^ on 15 mm Ø 24-well coverslips (500 μL/well) and treated to develop the NPC-phenotype. NPC-like fibroblasts were incubated with increasing concentrations of CDs (0.5, 1, 5, and 10 mM) for 48 h. At the end of the treatment, the medium was removed, and the cells were washed with PBS (once) and fixed with 4% PFA for 10 min. The fixing solution was removed, cells were washed with PBS (3 times) and stained with filipin (0.05 mg mL^−1^ in PBS) for 2 h at room temperature in the dark. Cells were then rinsed with PBS (3 times), mounted on glass slides and analyzed by DMI600 wide field fluorescence microscope (Leica microsystems, Wetzlar, Germany). Three images/condition were acquired (excitation 340-380 nm, emission 450-490 nm, 40x magnification). Filipin fluorescence intensity (FI) was obtained by LAS X software (Leica microsystems, Wetzlar, Germany) and divided by the number of cells/image. The value obtained from untreated NPC-fibroblasts was considered as 100%.

#### 4.3.7 Duration of the in vitro CD effect

L929 cells were seeded at 2.5 x 10^4^ mL^−1^ on 15 mm Ø 24-well coverslips (500 μL/well) and treated to develop the NPC-phenotype. NPC-like fibroblasts were incubated with the CDs (5 mM) for 48 h. The medium was then removed and replaced with CD-free complete medium. At different time points (24 or 48 h after CD treatment) the cells were stained with filipin and analyzed by fluorescence microscopy as described above. The value obtained from untreated fibroblasts was considered as 100%.

#### 4.3.8 CD pre-complexation experiments

CD-cholesterol complexes were prepared by adjusting a protocol described in dos Santos *et al*. [72]. Briefly, 1.5 mg of cholesterol were added to a 10 mL aqueous solution containing 10 mM of native or crosslinked HPβCDs and stirred for 5 days at 55 °C until complete cholesterol dissolution. The samples were collected and frozen overnight at −20 °C and then lyophilized for 2 days. L929 cells were seeded at 2.5 x 10^4^ mL^−1^ on 15 mm Ø 24-well coverslips (500 μL/well) and treated to develop the NPC-phenotype. NPC-like fibroblasts were incubated with CD-cholesterol complexes (obtained from monomeric or crosslinked HPβCDs). At the end of the treatment, the cells were stained with filipin and analyzed by fluorescence microscopy as previously described. The % of cholesterol has been obtained by the filipin FI divided by the number of cells and then normalized to CD-treated NPC-L929 cells (100%).

#### 4.3.9 CD uptake. Kinetics

L929 cells were seeded at 2.5 x 10^4^ mL^−1^ on 15 mm Ø 24-well coverslips (500 μL/well) and treated to develop the NPC-phenotype. NPC-like fibroblasts were incubated with 5 mM (based on HPβCD units) of fluorescently labeled monomeric, 19.3 kDa or 312.3 kDa HPβCDs up to 48 h. At selected time points (1, 2, 4, 8, 24 and 48 h after incubation) the medium was removed, and the cells were stained with filipin and analyzed by fluorescence microscopy as described above. In addition to filipin, RhB signal was acquired at excitation 525-550 nm, emission 585-640 nm. ***Inhibition of active processes*** - L929 cells were seeded at 2.5 x 10^4^ mL^−1^ on 15 mm Ø 24-well coverslips (500 μL/well) and treated to develop the NPC-phenotype. The cells were washed with complete medium and kept at 37 °C, 5% CO2 for 24 h. Then, NPC-like fibroblasts were either pre-incubated (30 min) at 4 °C or pre-treated (1 h) with adenosine triphosphate depletion solution (ATPd, 10 mM sodium azide and 6 mM 2-deoxy-D-glucose) at 37 °C. Afterwards, the cells were washed with complete medium and incubated with 5 mM of fluorescent monomeric, 19.3 kDa or 312.3 kDa HPβCDs for 8 h while keeping the temperature at 4 °C and 37 °C, respectively. At the end, the cells were stained with filipin and analyzed by fluorescence microscopy as described before. ***Confocal laser scanning microcopy*** - Samples from kinetics and inhibition of the active processes were analyzed by Fluoview 3000 (Olympus, Tokyo, Japan). One z-stack/condition was acquired (405, 488, and 594 nm lasers at 63x magnification using an oil-immersion objective) while the z-projections were elaborated using the ImageJ software (National Institute of Health, Bethesda, MD).

#### 4.3.10 Biodistribution and pharmacokinetics

Seven jugular vein-catheterized adult female Sprague Dawley rats were intravenously injected with 200 mg kg^−1^ of fluorescently labeled CD (either monomeric or 19.3-kDa HPβCD) and euthanized 12 h later. After overdose of xylazine and ketamine, animals were intracardially perfused with 4% PFA. Blood sampling was performed at 1, 2, 4, 8 and 12 h post CD injection, while main organs (cerebrum, cerebellum, spinal cord, lung, liver, kidney, spleen, pancreas and intestine) were collected after animal perfusion (*i.e*., after 12 h). Plasma was isolated by centrifugation (5000 x *g* for 5 min at 4 °C) and stored at −80 °C until CD quantification while organs were prepared following a protocol described by Polomska *et al*. [73]. Briefly, 0.1 g of tissue per 1 mL of PBS was homogenized with stainless steel beads of 5 mm (25–30 Hz for 5 min) by a Tissuelyser (Qiagen, Venlo, Netherlands). The homogenates were centrifuged at 10,000 x *g* for 10 min and the supernatant was collected and stored at −80 °C until CD quantification. All the samples were then analyzed by Infinite M200 (Tecan, Zürich, Switzerland), to quantify fluorescent CDs (543 nm excitation/580 nm emission, 100% gain, integration time 20 μs, number of flashes: 20). Calibration curves were obtained for both CDs in blood and organs of untreated rats. In order to evaluate the penetration of the CDs in the organs’ parenchyma, small sections of all were collected before homogenization. These sections were sliced and stored at −80 °C until further manipulation. Immunofluorescence was performed on 10-μm lung, liver, kidney and brain serial sections, treated with cold methanol for 10 min at −20 °C, washed 3 times in PBS, stained for nucleus with Hoechst (1: 1000 in PBS) for 20 min at RT and then mounted on glass slides. At least 3 pictures/section were acquired at 10x (lung, liver and kidney) and 40x (brain) magnification by DMI600 wide field fluorescence microscope (Leica microsystems, Wetzlar, Germany). Hoechst signal (cell nucleus) was acquired at excitation 340-380 and emission 450-490 channels and RhB (crosslinked CD) was acquired at excitation 542-582 and emission 604-644 channels. In the pharmacokinetic study, CD concentration in the plasma was expressed in μg mL^−1^ while t_1/2_, C_max_, AUC 0-t, CL, Vd and MRT were calculated using the Excel-based function PK Solver 2.0 plugin (Microsoft Corporation, Redmond, WA) by setting a non-compartmental analysis after i.v. bolus input. CD concentration in the organs was expressed as % of injected dose (ID) per g of tissue. Additional 4 rats were injected with 200 mg kg^−1^ of fluorescent 19.3 kDa HPβCD and euthanized 24 h later. Blood samples were collected at 24 h post CD injection and treated as described above.

#### 4.3.11 Therapeutic efficacy in vivo

The *in vivo* was performed in a blinded fashion. The person administering the CDs and following the life span of the animals did not know which groups were receiving the monomeric HPβCD and which groups were receiving the crosslinked HPβCD. There were four groups (*n*=6 for the lower dose of monomeric HPβCD*, n*=5 for the higher dose of monomeric HPβCD and *n*=4 for both doses of crosslinked HPβCD). The Npc1 mice received intraperitoneal injection on postnatal day 7 (P7) with either crosslinked HPβCD or monomeric HPβCD at the dose of 1333 or 4000 mg kg^−1^. The injections were performed with 7 or 20% (w/v) CD in phosphate buffered saline, respectively. Survival is expressed as % of the cumulative number of dead animals *vs*. the initial number of animals per group.

#### 4.3.12 Magnetic resonance imaging-guided low intensity-pulsed focused ultrasound

***Microbubbles***- Microbubbles were prepared adapting the protocol developed by Feshitan *et al*. [74]. Briefly, a lipid suspension of 90 mol% DSPE, 5 mol% DSPE PEG 2 kDa and 5 mol% DSPE PEG 5 kDa was prepared at 2 mg/mL in 100 mL PBS. The suspension was degassed, preheated to 80 °C and then sonicated at low power (3 W for 10 sec). Perfluorobutane gas was introduced by flowing it over the surface of the solution and higher power sonication (33 W for 10 sec) was applied. The suspension was collected into 30 mL syringes (Tyco Healthcare, Mansfield, MA), which were used as the flotation columns. Centrifugation (300 x *g* for 5 min) was performed to collect all microbubbles from the suspension into a cake and stored at 4 °C until their use (within 24 h). The median diameter of the microbubbles was 1.556 μm and the suspension was diluted (57 μL of microbubbles in 543 μL of PBS) before administration. ***BBB opening setting***- Four jugular vein-catheterized adult female Sprague Dawley rats were anesthetized with isoflurane (2.5% in a mixture 4:1 air:oxygen) and placed into the animal support of the rodent FUS system (IGT, Pessac, France). Their skull region was shaved, and the body temperature of the animals was kept constant (36-37 °C) using a hot water circuit integrated into the animal support. T1-weighted anatomical MR images were acquired on a Bruker BioSpec 470/30 small animal MR system (Bruker, Billerica, MA) operating at 7.0 T and used to set the position of the 1.5 MHz FUS transducer on the head of the animals. The rats were i.v. injected with 2.4 mL kg^−1^ of microbubbles (600 μL/min injection speed) 30 s before the beginning of the FUS sequence, which lasted for 3 min (Length: 5-10 ms pulse; Sequence: 1 Hz pulse repetition frequency; Pressure *in situ:* 0.4-0.5 mPa) and was adapted from a protocol described by Papachristodoulou *et al*. [59]. Twenty min after FUS sequence, rats were i.v. injected with 75 μL/animal of Gadolinium-DOTA (Guerbet, Paris, France) and T1-weighted MR images were acquired to confirm BBB opening. Then, one rat was i.v. injected with Evan’s blue and euthanized by an overdose of xylazine and ketamine, the brain was collected and analyzed to confirm BBB opening. ***Combination of MRIg-FUS and CDs***- Eight jugular vein-catheterized adult female Sprague Dawley rats were prepared as in 4.3.12. The rats were first i.v. injected with microbubbles 30 s before the beginning of the FUS sequence, and then with 200 mg kg^−1^ of fluorescently labeled crosslinked CD. The FUS sequence lasted for 3 min and BBB opening confirmed with T1-weighted contrast-enhanced MRI. Animals were euthanized 8 h later with an overdose of xylazine and ketamine and intracardially perfused with 4% PFA. The brains were collected, stored at −80 °C and sliced to be further analyzed. Immunofluorescence was performed on 15-μm brain serial sections, treated with cold methanol for 10 min at −20 °C, washed 3 times in PBS and incubated with BSA 1% and Triton 100x 0.5% (both in PBS) for 30 min at RT. Sections were then incubated either with rabbit anti-Olig2 (1:100 in PBS) or with rabbit anti-GFAP (1:200 in PBS) primary antibodies overnight at 4 °C and stained for oligodendrocytes and astrocytes, respectively, with Alexa goat anti-rabbit Alexa 488 secondary antibody (1:250 in PBS) for 90 min at RT. Sections were ultimately stained for nucleus with Hoechst (1:1000 in PBS) for 20 min at RT and then mounted on glass slides. At least 3 pictures/section were acquired at 40x and 63x magnification by DMI600 wide field fluorescence microscope (Leica microsystems, Wetzlar, Germany). Hoechst signal (cell nucleus) was acquired at excitation 340-380 and emission 450-490, Alexa 488 channels (either astrocytes or oligodendrocytes) was acquired at excitation 460-500 and emission 512-542 channels, and RhB (crosslinked CD) was acquired at excitation 542-582 and emission 604-644 channels.

#### 4.3.13 Data Analysis

Experiments were performed at least 3 times. *n* refers to the number of independent experiments. The statistical analysis (statistical significance p < 0.05) and the graphs were made by GraphPad Prism 8.0. Data are expressed as mean + standard deviation (SD). The type of statistical test and post-hoc multiple comparison is specified in the caption of each figure and was selected according to the number of replicates/groups and the type of comparison. and the suggestion of the software. Outliers were identified and excluded by GraphPad QuickCalcs.

#### 4.3.14 Data Availability

Main data are reported in the article. Supporting information (*i.e*., representative ^1^H NMR spectra, images and controls for all experiments) are provided in a separate file. Any other raw data are available upon request.

## Acknowledgements

A special acknowledgement to Mr. Christoph Poincilit for his constant personal and technical support. The authors thank ScopeM (ETH Zurich) for the use of their microscope platform. Dave Smith and Claire Fletcher (University of Oxford) are gratefully acknowledged for breeding the transgenic Npc1 mice. The authors thank Prof. Yanik M. Fatih and Dr. Wolfger von der Behrens (ETH Zurich and University of Zürich) for the administrative support with the animal license, the use of their rat husbandry and MRI platform. Dr. Anastasia Spyrogianni Roveri and Dr. Salvatore Cinquerrui (ETH Zürich) are acknowledged for the review of the original paper draft. The authors thank Paul Johnson (ETH Zürich and University of Zürich) to produce the microbubbles used in the MRIg-FUS experiments. This project was financially supported by the Novartis Foundation for Medical-Biological Research, Vontobel foundation, Carigest SA and NPSuisse. Hsintsung Chen is a recipient of an Oxford-Taiwan Graduate Scholarship jointly funded by the University of Oxford and the Ministry of Education of the Republic of China. The University’s share of this scholarship is jointly funded by two charitable organisations, the Niemann-Pick Research Foundation (NPRF) & Niemann-Pick UK (NPUK). Frances Platt is a Royal Society Wolfson Research Merit Award holder and a Welcome Trust Investigator in Science.

## Conflict of Interest

The authors declare no conflict.

## Author Contributions

Dr. Dario Carradori co-designed the study, performed the experiments, analyzed and interpreted the data, and wrote the original draft of the manuscript. Hsintsung Chen and Prof. Frances Platt designed, performed and assisted with the interpretation of the *in vivo* study on the NPC mouse model and funded this element of the project (NPUK and NPRF). Beat Werner and Aagam Shah assisted with the FUS study on healthy rats. Chiara Leonardi assisted with the *in vitro* mechanistic studies of HPβCDs. Mattia Usuelli and Prof. Raffaele Mezzenga assisted with the execution and interpretation of SLS measurements. Prof. Jean-Christophe Leroux, co-designed the study, obtained funding, supervised the project, and guided experimental design, data interpretation and manuscript preparation. All authors reviewed the manuscript.

## Notes

### Competing Interest Statement

The authors have declared no competing interest.

