## Supplementary Info for "Elucidating the mechanism of cyclodextrins in the treatment of Niemann-Pick Disease Type C using crosslinked 2-hydroxypropyl-β-cyclodextrin"

***1. Representative ^1^H NMR spectra of crosslinked HPβCDs.***

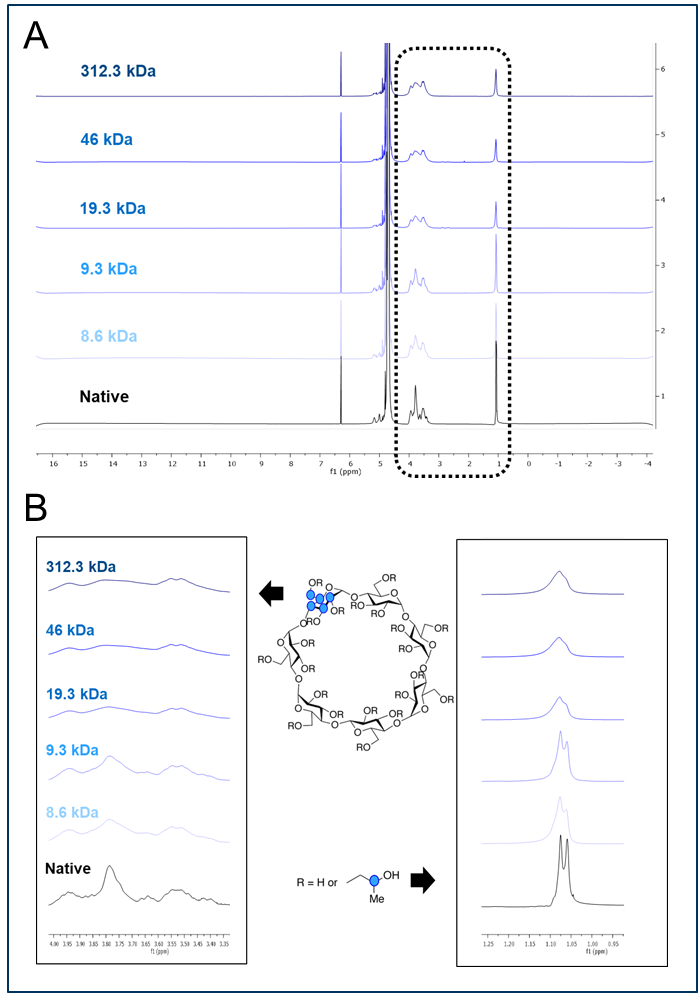

**Figure 1S. ^1^H NMR spectra of crosslinked** **HPβCDs.** Ten mg of each CD were dissolved in 600 µL of deuterium oxide containing 35 mM of maleic acid (internal standard). Samples were analyzed with a ^1^H NMR 400 MHz spectrometer. **A**, full spectra. **B**, details showing the modified regions after cross-linking.

***2. Representative SLS analysis of crosslinked HPβCDs.***

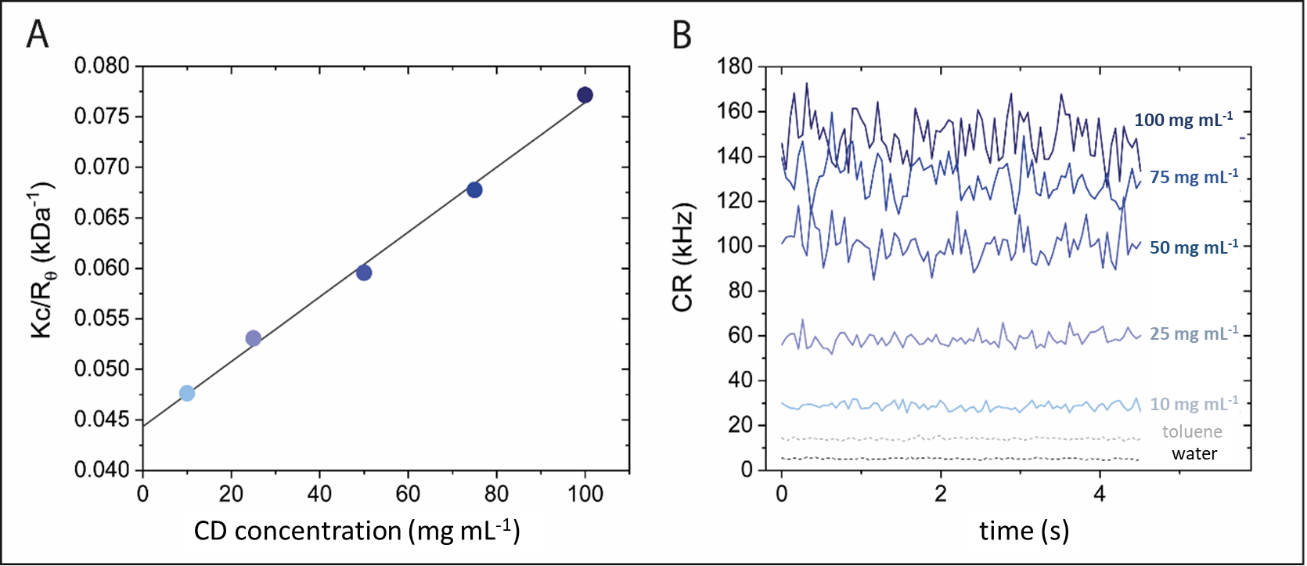

**Figure 2S. SLS analysis of the 19.3 kDa crosslinked HPβCD.** **A**) Debye Plot and **B**) raw count rate (CR) at different CD concentrations. Kc corresponds to the optical constant (obtained from the laser wavelength and the refractive index of solvent and sample) and R_θ_ is the Rayleigh ratio of the sample. Toluene and water were used as internal controls.

***3. Cytotoxic profile of normal fibroblasts (L929) after incubation with crosslinked HPβCDs.***

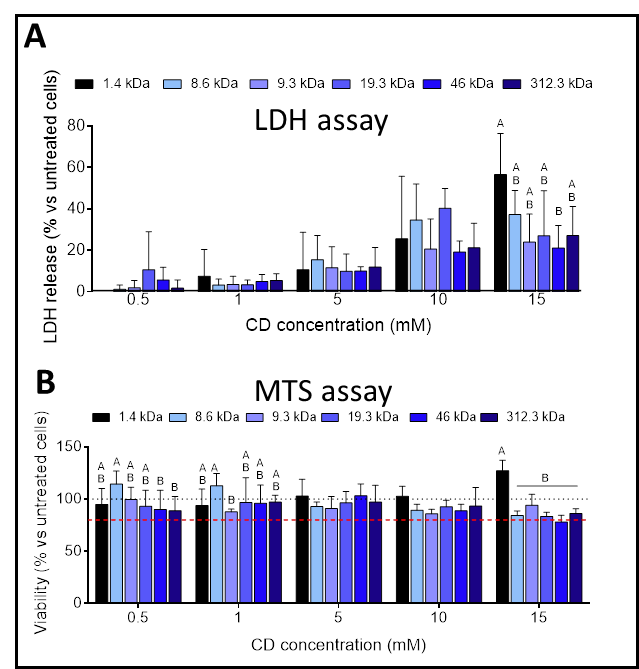

MTS assay

LDH assay

**Figure 3S. CD effect on normal fibroblasts.** CD treatment was performed at 37 °C in CO_2_ 5% for 48 h. **A**) LDH release and **B**) cell viability after CD-treatment. The % of LDH release and the % of cell viability were normalized to untreated NPC-L929 cells (100%). Two-way ANOVA with multiple comparisons (Tukey’s post-hoc test). Mean + SD (*n*=3). Different letters indicate a statistically significant difference (p < 0.05).

***4. CD cellular uptake***

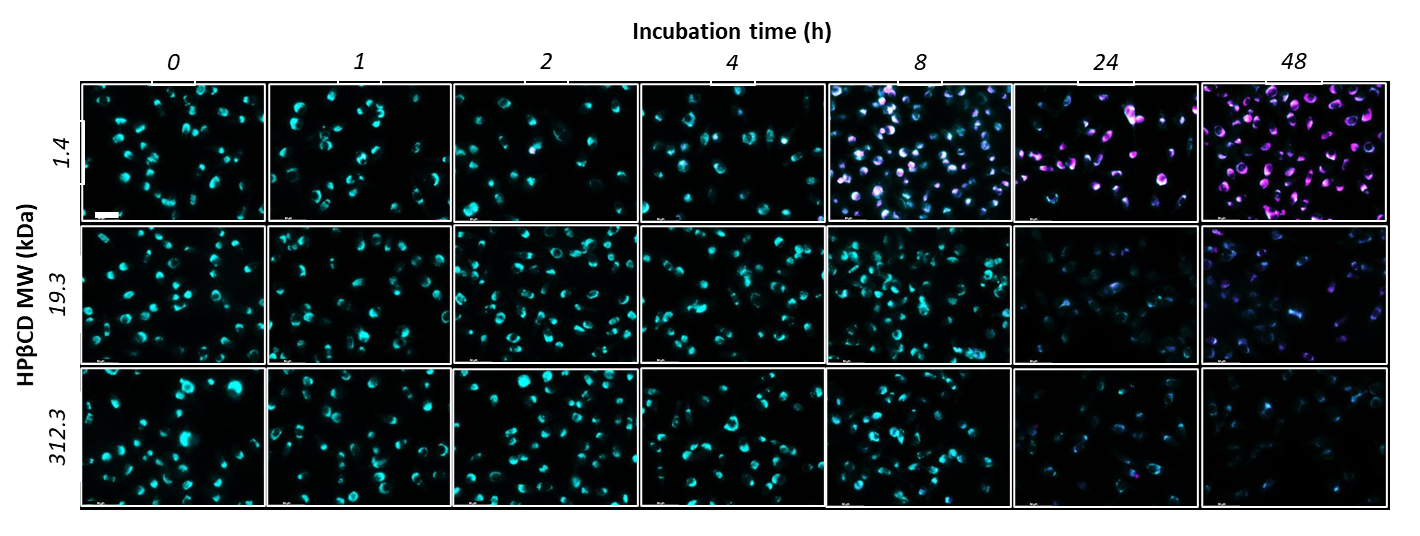

**Figure 4S. CD cellular uptake at different time-points.** Representative pictures of NPC-L929 cells incubated for different times (1, 2, 4, 8, 24, and 48 h) with 5 mM fluorescent CDs (monomeric, 19.3 or 312.3 kDa HPβCD). Cyan corresponds to filipin fluorescence (*i.e.,* intracellular cholesterol accumulation) and pink corresponds to RhB (*i.e.,* fluorescent CDs). Scale bar: 50 µm.

***5. Orthogonal sections***

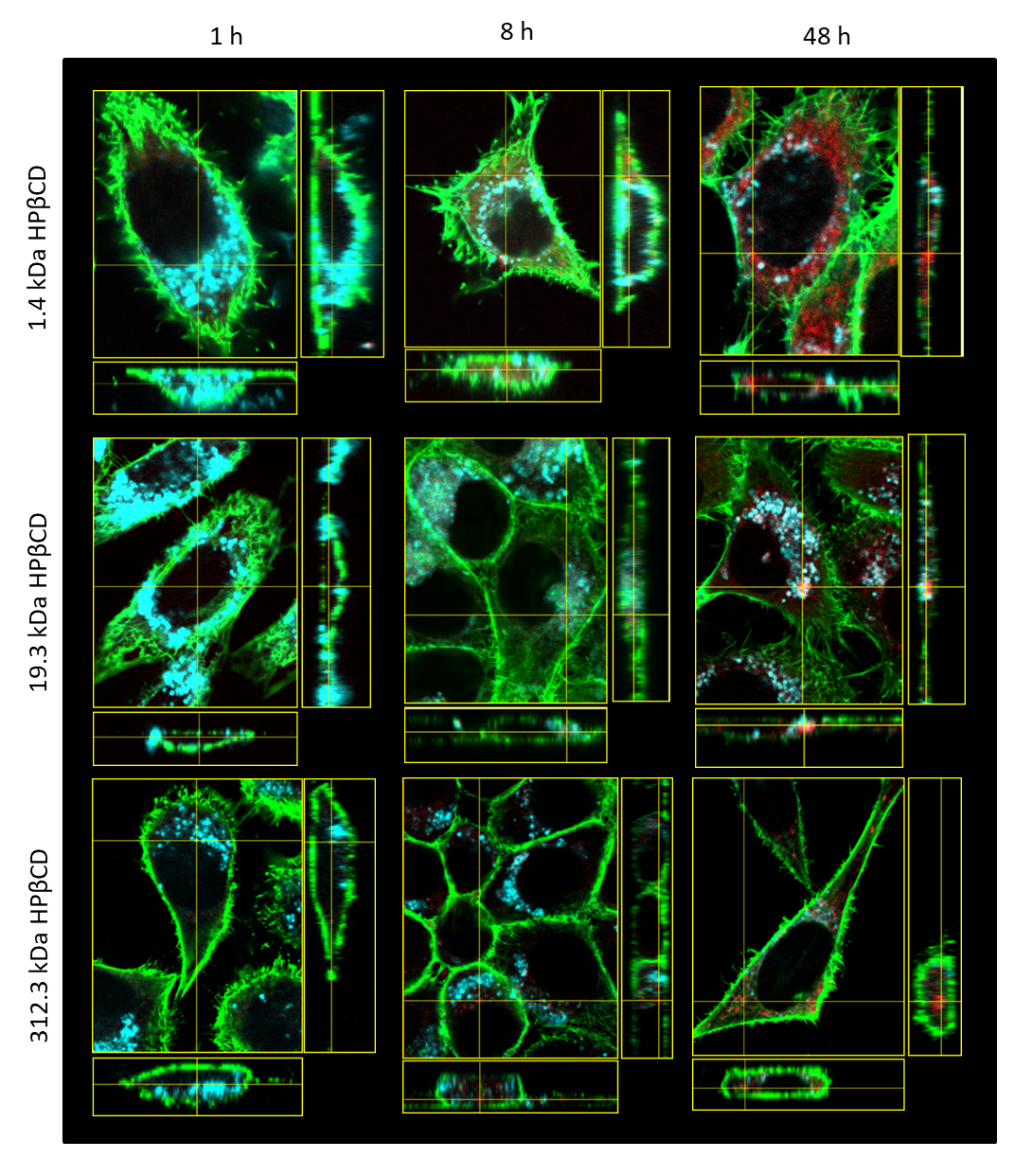

**Figure 5S. CD localization at different time-points.** Representative orthogonal section pictures of NPC-L929 cells incubated for different times (1, 8, and 48 h) with 5 mM fluorescent CDs (monomeric, 19.3 or 312.3 kDa HPβCD). Cyan corresponds to filipin fluorescence (*i.e.*, intracellular cholesterol accumulation), green is phalloidin-488 (*i.e.,* cytoskeleton) and red corresponds to RhB (*i.e.,* fluorescent CDs).

***6. Magnetic resonance imaging-guided low intensity-pulsed focused ultrasound***

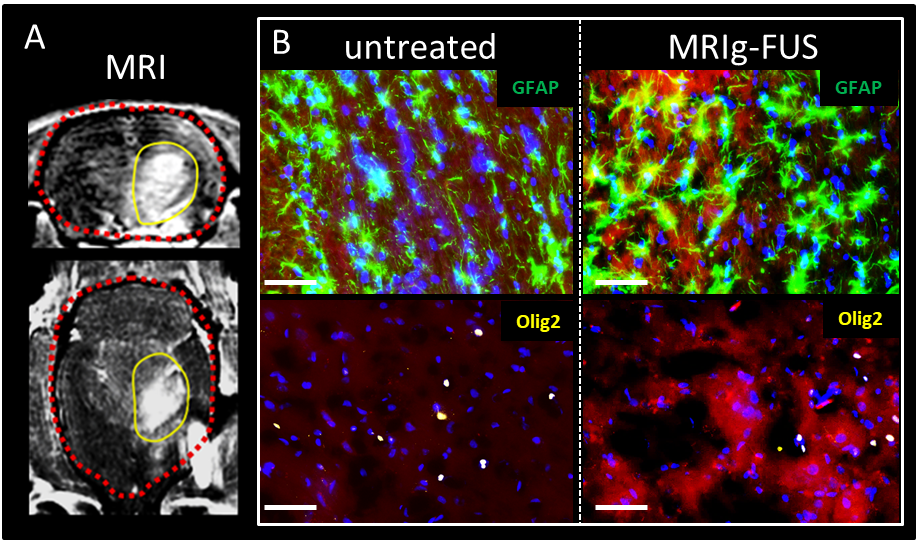

**Figure 6S. Brain diffusion of monomeric CD after MRIg-FUS.** **A**) T1-weighted MRI of gadolinium extravasation (clear grey) after MRIg-FUS. **B**) Representative pictures showing the localization of 1.4 kDa HPβCD compared to the astrocytes (green) and oligodendrocytes (yellow) with and without MRIg-FUS. Blue corresponds to Hoechst (*i.e*., cell nuclei), green corresponds to GFAP (*i.e*., astrocytes), yellow corresponds to Olig2 (*i.e*., oligodendrocytes), red corresponds to RhB (*i.e*., fluorescent CD), Scale bar: 50 µm.
